# Glypican-1 upregulation elicited in response to a cell-impermeable kinase inhibitor and its overexpression enhance HIV-1 infection

**DOI:** 10.1101/2025.10.28.685002

**Authors:** Kelsey Vinzant, Natalia Cheshenko, Heng Pan, Ronald Cutler, Joshua N. Buckler, Andreas Luxenburger, Lawrence D. Harris, Jeffrey R. Johnson, Steven C. Almo, Betsy C. Herold

**Author notes:** Corresponding author (BCH).

## Abstract

Studies of herpes simplex virus (HSV) entry uncovered a previously unappreciated “outside-in” signaling pathway whereby activation of the calcium (Ca^2+^) responsive enzyme phospholipid scramblase 1 (PLSCR1), which is known to trigger the bidirectional movement of phosphatidylserines (PS) between the inner and outer leaflet of the plasma membrane, also induces the translocation and subsequent extracellular activation of intracellular proteins, including Akt. We hypothesized that HIV-1, which has been shown to elicit scramblase TMEM16F-mediated PS externalization, may trigger a similar “outside-in” signaling cascade involving exofacial kinase activity to promote its entry into CD4+ T cells. To study this process, we utilized a cell impermeable staurosporine analogue, alkyl-CIMSS, which is a broadly active kinase inhibitor that blocks HSV-induced exofacial Akt phosphorylation and HSV infection. Using multiple cell types including TZM-bl, Jurkat T cells, and human peripheral blood mononuclear cells (PBMCs), we show that, in contrast to the effects on HSV, treatment of cells with alkyl-CIMSS enhances HIV-1 infection post-entry that is not dependent on TMEM16F. To identify potential biological processes that are responsive to alkyl-CIMSS, we performed bulk RNA-sequencing and whole cell proteomics and found that alkyl-CIMSS treatment of cells robustly upregulates the cell surface density of the proteoglycan glypican-1 (GPC1). Lentiviral delivery of GPC1 overexpression and shRNA knockdown constructs reveal that the presence and absence of GPC1 independently of alkyl-CIMSS treatment significantly impact HIV-1 infection, with the effect on infection corresponding to GPC1 expression. Further, we demonstrate that the influence of GPC1 on HIV-1 infection is in part mediated by TGF-β signaling. Collectively, these findings implicate a cell surface protein susceptible to alkyl-CIMSS in restricting HIV-1 infection and identify GPC1 as a novel modulator of HIV-1 infection.

**Author Summary:** We utilized a cell-impermeable pan-kinase tool compound, alkyl-CIMSS, to identify cell surface molecules that might be involved in viral infection. Treatment of CD4+ T cells with alkyl-CIMSS increased the expression of the cell-surface protein glypican-1, which led to an increase in early HIV reverse transcriptase products and promoted viral infection. Conversely, alkyl-CIMSS inhibited herpes simplex virus entry and infection. These findings illustrate that viruses interact with exofacial cell membrane molecules differently to promote or impede infection.

## Introduction

Viruses usurp host cell machinery to promote their entry and replication [1–3]. Prior investigations of herpes simplex virus (HSV) entry into human epithelial cells uncovered a previously unappreciated paradigm associated with activation of phospholipid scramblase 1 (PLSCR1), a calcium (Ca^2+^)-responsive enzyme known to catalyze the movement of phosphatidylserine lipids (PS) between the inner (cytofacial) and outer (exofacial) leaflet of the plasma membrane (PM) [4]. Surprisingly, we found that the movement of PS to the exofacial leaflet in response to HSV was associated with the concomitant translocation of intracellular proteins, including the master kinases Akt and phosphoinositide-dependent kinase 1 (PDPK1), as well as other signaling molecules [5, 6]. Phosphorylation of exofacial Akt and PDPK1 triggered downstream intracellular signaling responses (“outside-in” signaling), which culminated in fusion of the HSV envelope and the PM, release of viral capsids into the cytoplasm and their transport to the nuclear pore to initiate viral replication.

To further study this “outside-in” signaling pathway activated by HSV, we previously designed a <u>c</u>ell-<u>im</u>permeable analog of the broadly active kinase inhibitor<u>s</u>tauro<u>s</u>porine (abbreviated CIMSS) as a tool compound. We confirmed the cell-impermeability of CIMSS and demonstrated that it blocked the phosphorylation of exofacial kinases including Akt and prevented HSV entry but did not inhibit the phosphorylation of intracellular Akt [6]. For example, activation of cytoplasmic Akt in response to insulin binding to the insulin receptor was not perturbed. Vesicular stomatitis virus pseudotyped with SARS-CoV-2 spike (VSV-S) and native SARS-CoV-2, but not native VSV, activated a similar exofacial signaling cascade and were susceptible to CIMSS inhibition.

HIV-1 initiates infection by binding to CD4 and then to either of the CCR5 or CXCR4 coreceptors, which are chemokine receptors that trigger Ca^2+^ fluxes [7]. The increase in intracellular Ca^2+^ in response to HIV-1 coreceptor engagement was previously shown to activate a different transmembrane phospholipid scramblase, TMEM16F, a member of the family of Ca^2+^-activated chloride channels and scramblases (CaCCs) [8]. Pharmacological inhibition or knockdown of TMEM16F blocked PS translocation and inhibited HIV-1 infection [8]. We hypothesized that HIV-mediated activation of TMEM16F might also trigger the exofacial translocation of Akt and/or other intracellular kinases and render HIV-1 susceptible to inhibition by CIMSS. To test this hypothesis, we compared the susceptibility of HIV-1 and HSV to a modestly modified alkyl version of the original acyl-CIMSS in model cell culture systems and primary peripheral blood mononuclear cells (PBMCs). Consistent with our earlier studies, HSV induced exofacial movement of both PS and Akt in HeLa cells and infection was inhibited by alkyl-CIMSS. In contrast, HIV-1 triggered exofacial movement of PS but not Akt. Moreover, not only did alkyl-CIMSS fail to inhibit HIV-1 infection, but surprisingly, treatment with alkyl-CIMSS led to an increase in HIV-1 infection and the increased production of early reverse transcriptase products. Transcriptomic and proteomic studies showed that alkyl-CIMSS treatment altered the cell transcriptome and proteome, in particular upregulating glypican-1 (GPC1). Lentiviral introduced overexpression or knockdown of GPC1 demonstrated that increased levels of GPC1 promote HIV-1 infection. Further, increased infection by either alkyl-CIMSS or genetic manipulation of GPC1 could be reduced by chemical inhibition of TGF-β signaling.

These findings illustrate that viruses interact with host extracellular cell membrane molecules differently to promote infection.

## Results

### A cell-impermeable analog of staurosporine enhances HIV-1 infection

Inhibition of exofacial kinases with acyl-CIMSS, a nonselective cell impermeable modified staurosporine pan-kinase inhibitor, inhibited HSV-2 and SARS-CoV-2 entry into epithelial cells downstream of PLSCR1 activation by blocking phosphorylation of exofacial kinases including Akt [6]. We hypothesized that HIV-1, which activates a different Ca^2+^ dependent scramblase, TMEM16F [8], might also be susceptible to CIMSS inhibition. We modified the previously described acyl-CIMSS by attaching the impermeabilizing sulfonate group via 4’-amine-alkylation (**S1 Fig. A and S2 Fig.**) rather than through an amide bond, as the amide linkage has been hypothesized to decrease binding by loss of hydrogen bonding [9]. Additionally, an alkyl linkage would be expected to provide greater stability than an amide bond, but both were found to be otherwise functionally identical and alkyl-CIMSS and retained the cell impermeability and kinase inhibition properties of the original CIMSS (**S1 Table**). Consistent with its cell impermeability, alkyl-CIMSS had no discernible effects on cell growth or viability when TZM-bl cells, a HeLa cell line engineered to express CD4, CCR5, CXCR4, and a luciferase reporter under the control of the HIV-1 long terminal repeat *(ltr*), Jurkat-CCR5 CD4+ T cells, or primary human CD4+ T cells were incubated with the drug for 24 hours (**S1 Fig. B**). Moreover, unlike unmodified staurosporine, alkyl-CIMSS did not trigger apoptosis as measured by caspase activation (**S1 Fig. C**).

At the previously established 50% inhibitory dose of the acyl-CIMSS for HSV entry into epithelial cells, 10 μM alkyl-CIMSS reduced HSV-2 infection of TZM-bl, Jurkat-CCR5 and primary CD4+ T cells, by a mean of 63±0.010%, 49±0.080% and 43±0.080%, respectively, relative to vehicle (DMSO)-treated cells. Infection was assessed by quantifying intracellular ICP0 expression, an immediate early HSV gene, 4 hours post viral exposure (**Fig 1A**). In surprising contrast, not only did alkyl-CIMSS fail to inhibit HIV-1 infection, but it significantly increased both HIV-1_BaL_ (R5-utilizing strain) and HIV-1_IIIB_ (CXCR4-utilizing strain) infection of TZM-bl cells in a dose dependent manner relative to cells treated with DMSO control (**Fig 1B and C**). TAK-779, a CCR5 antagonist, and AMD3100, a CXCR4 antagonist, were included as negative controls. Based on the dose response curve with TZM-bl cells, we selected 50 µM alkyl-CIMSS for subsequent experiments, as this concentration enhanced HIV-1_BaL_ infection 4.8 ± 2.3-fold. Given that R5-tropic viruses are the predominant strains responsible for primary HIV-1 infections [10], we focused our subsequent studies on R5 viruses. Consistent with the TZM-bl data, alkyl-CIMSS significantly increased HIV-1_BaL_ infection of Jurkat-CCR5 (**Fig 1D-G**). The increase in infection of Jurkat-CCR5 cells was observed both by directly quantifying p24 expression by flow cytometry or, indirectly, by collecting culture supernatants and quantifying viral yields in a TZM-bl luciferase assay. We also observed an increase in p24 released into the media and by quantifying viral yields when primary human PBMCs were infected with HIV-1_BaL_ in the presence of alkyl-CIMSS, although the response was smaller and more variable than observed with cell lines (**Fig 1H and I)**.

**Fig 1:**
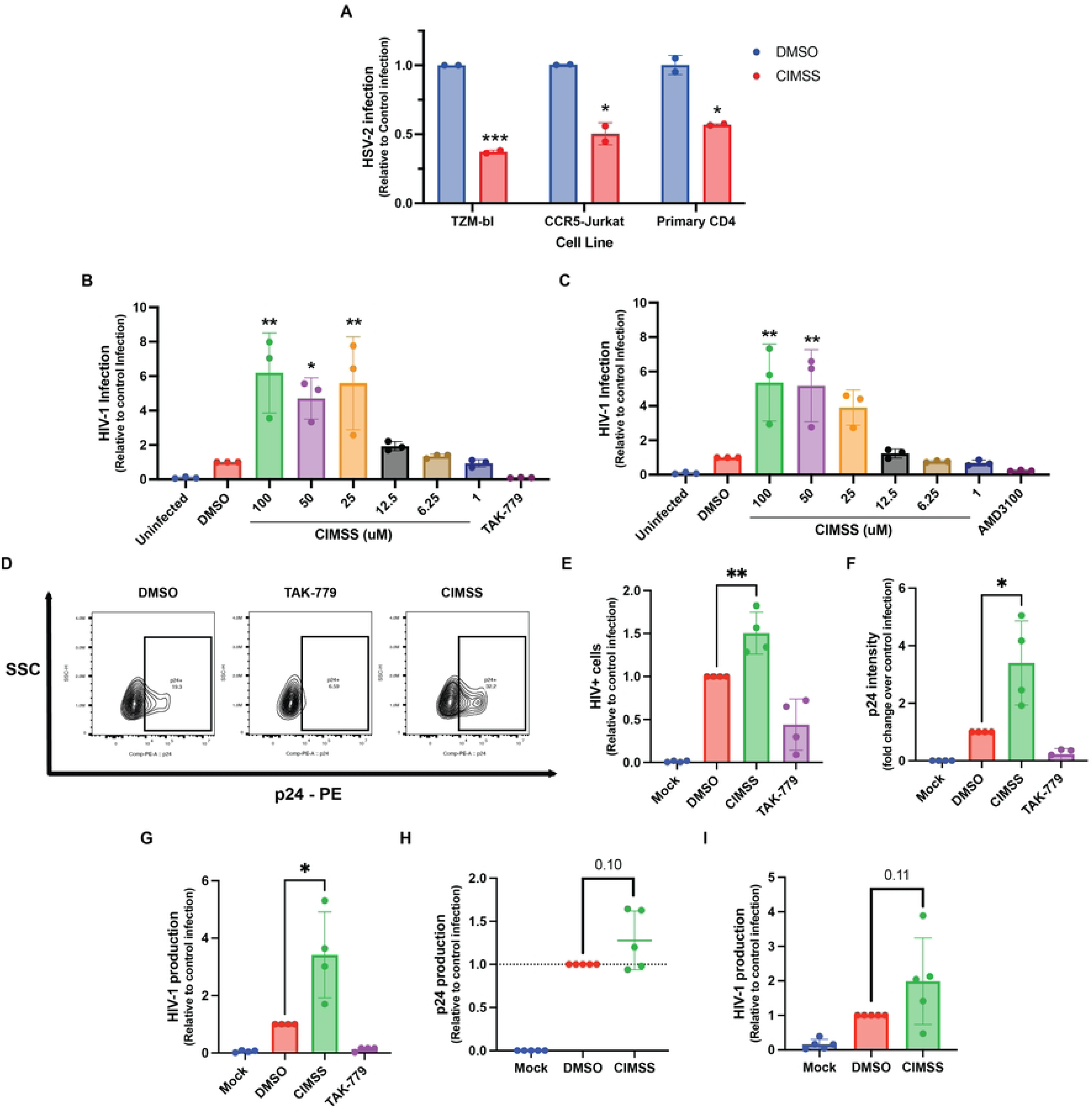
Alkyl-CIMSS inhibits HSV-2 but enhances HIV-1 infection. (A) The indicated cells were treated with 10 μM alkyl-CIMSS or DMSO vehicle control and then infected with HSV-2(G) (10 pfu/cell) for 4 hours. Infection was monitored quantifying ICP0 expression in isolated RNA (n = 2 independent experiments each conducted in duplicate per cell type). (B and C) TZM-bl cells were treated with the indicated concentrations of alkyl-CIMSS, DMSO vehicle control, 10 μM TAK-779 (R5) or 10 μM AMD3100 (X4) and then infected with 1.5 ng p24 of HIV-1_BaL_ (R5) (B) or HIV-1_iiiB_ (X4) (C). Infection was quantified by lysing cells for luciferase expression after 48 h incubation (n=3 independent experiments each conducted in duplicate for each viral strain). (D-F) Jurkat-CCR5 cells were treated with 50μM CIMSS, DMSO vehicle control or 10 μM TAK-779 and infected with 5 ng p24 HIV-1_BaL_. At 72 hpi, cells were fixed and permeabilized for p24 and stained for cell viability, A representative flow plot is shown in D, percentage p24% cells in E and mean fluorescence intensity in F (n=4 independent experiments each conducted in duplicate). (G) Viral yields from infected Jurkat-CCR5 cells from (D) were quantified by TZM-bl luciferase assay. (H) Whole PBMCs were treated with the indicated compound and infected with HIV-1_BaL_ for 5 days. Viral production was quantified by measuring p24 by ELISA and (I) yields by TZM-bl luciferase assay in culture supernatants (n = 4). Results are presented as mean ± SD relative to cells infected in the presence of the DMSO vehicle control and compared by one-way ANOVA (B, C) or unpaired t-test (A, E-I); *P < 0.05, **P < 0.01, ***P <0.001, ****P <0.0001.

### HIV-1 triggers exofacial movement of phosphatidylserine lipids but not Akt

Exofacial Akt phosphorylation was previously identified to be important for HSV and SARS-CoV-2 entry and was inhibited by treatment with acyl-CIMSS [6]. As alkyl-CIMSS did not inhibit but enhanced HIV-1 infection, we hypothesized that HIV-1 may not trigger translocation of Akt to the outer leaflet of the plasma membrane. To directly test this notion, we treated cells with HSV-2 or HIV-1 and assayed for exofacial PS and Akt. Both HSV-2 and HIV-1 triggered a significant increase in exofacial PS within 30 minutes of exposure (**Fig 2A**), but only HSV-2 exposure resulted in detectable exofacial Akt as assayed by flow of non-permeabilized TZM-bl cells (**Figs 2B and C).** Permeabilized cells were included as a positive control for detection of intracellular Akt.

**Fig 2:**
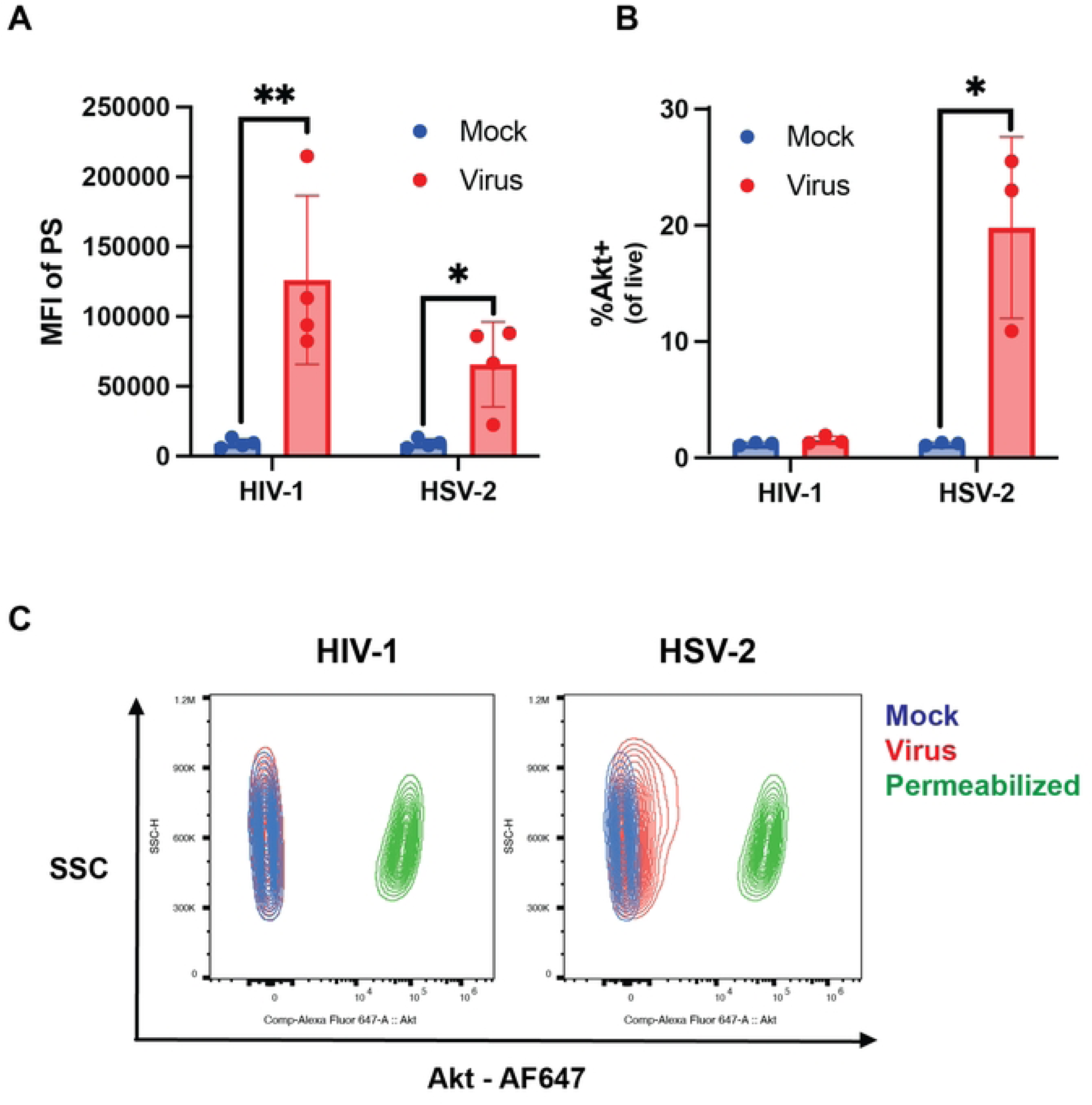
HIV and HSV trigger exofacial translocation of phosphatidylserines but only HSV increases exofacial Akt. (A) TZM-bl cells were treated with HIV-1 or HSV-2 for 30 minutes and stained for cell viability and phosphatidylserines (PS). Results are presented as the mean fluorescence intensity (MFI) of PS after gating on the live cell population (n=4 independent experiments). (B). Cells were infected as in A and the percentage of nonpermeabilized live cells expressing Akt quantified by flow (n=3 independent experiments). (C) Representative contour plots showing changes in Akt staining following virus treatment (blue: mock, red: virus treated, green: permeabilized). Results are shown as mean ± SD and compared by unpaired t-test; *P < 0.05, **P < 0.01.

Since both HIV-1 and HSV trigger an increase in exofacial PS but only HSV elicits translocation of Akt to the outer leaflet of the PM, we hypothesized that the difference may be explained, in part, by the findings that HSV activates PLSCR1 [5] whereas HIV-1 has been previously shown to activate TMEM16F [8]. To confirm that HIV-1 activates TMEM16F, we tested the effects of the pharmacological CaCCinh-A01 (A01) inhibitor, which blocks TMEM16F scramblase activity (**Fig 3A**) [11, 12]. We also engineered a Jurkat-CCR5 cell line in which TMEM16F was knocked out (KO) using CRISPR technology. The KO was confirmed by assessing gene and protein expression by RT-qPCR and flow cytometry, respectively (**Fig 3B**). Treatment of Jurkat-CCR5 cells with A01 (**Fig 3A**) or exposure of the TMEM16F KO cells to HIV-1 (5 ng p24) resulted in a significant reduction in HIV-1 infection as measured by the percentage of cells expressing viral p24 at 72 hours but did not block HSV infection, which was quantified using a reporter virus expressing mCherry linked to the HSV capsid protein, VP26, 4 hours post exposure (**Fig 3C**). In addition, there was significantly less PS detected exofacially by flow cytometry following HIV-1 exposure of the TMEM16F KO compared to the CRISPR control cells (**Fig 3D**).

**Fig 3:**
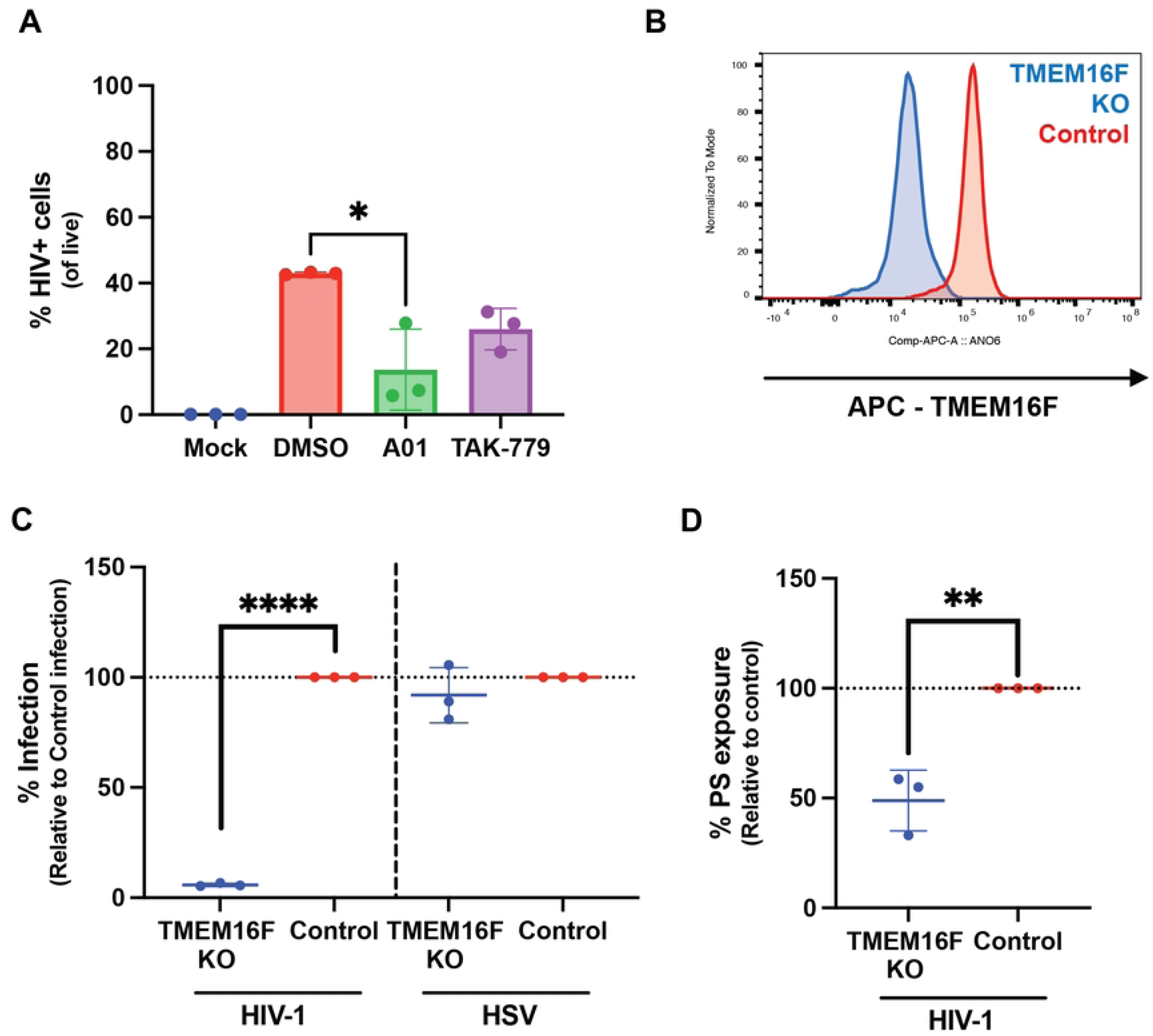
HIV but not HSV infection is reduced in the absence of TMEM16F. (A) Jurkat-CCR5 cells were treated with 10 μM TAK-779, or 60 μM CaCCinh-A01 (A01) or DMS0 vehicle control before infection with 5 ng p24 HIV-1_BaL_. Cells were fixed and permeabilized for p24 and viability staining after 72 h incubation (n=3 independent experiments each conducted in duplicate). Results are presented as percent of live cells. (B) Representative histogram showing TMEM16F expression in TMEM16F knockout (KO) (blue) and CRISPR control cells (red). (C) TMEM16F KO or control cells were infected with HIV-1 or HSV-VP26-mCherry and infection quantified by assessing the percentage of p24 or mCherry positive cells by flow cytometry 72- or 4-hours post-infection, respectively. Results are presented as the percent of HIV or HSV+ cells relative to control cells (n=3 independent experiments each conducted in duplicate). (D) TMEM16F KO or control cells were exposed to HIV-1_BaL_ for 30 minutes and then stained for exofacial PS by flow cytometry in nonpermeabilized cells. Results are presented as the MFI of PS on HIV-exposed cells normalized to the virus exposed control cells (n=3 independent experiments each conducted in duplicate). Results are shown as mean ± SD and compared by unpaired t-test; *P < 0.05, **P < 0.01, ****P <0.0001.

### Alkyl-CIMSS acts on the cell to promote expression of reverse transcriptase products

The observation that alkyl-CIMSS enhanced HIV-1 infection despite the absence of any detectable exofacial Akt suggests a mechanism distinct from that which is involved in the inhibition of HSV infection. To explore how alkyl-CIMSS enhances HIV-1 infection, we tested whether the compound targets a molecule accessible on the cell surface prior to or only following HIV-1 exposure or whether it directly interacts with components of the viral particle. TZM-bl cells were pretreated with alkyl-CIMSS (or control buffer) for one hour and then washed extensively prior to HIV-1 infection (1.5 ng p24), or concentrated virus (40x) was pretreated with the compound (or control buffer) for one hour and then diluted to a final inactive concentration of 1μM alkyl-CIMSS before infecting cells. As a positive control in both assays, the cells were exposed simultaneously to virus and alkyl-CIMSS. Enhancement of HIV-1 infection persisted if cells were pretreated with alkyl-CIMSS and then washed prior to infection (**Fig 4A**) but the effect observed with simultaneous addition of drug and virus was reduced when virus was pretreated with the drug and diluted (**Fig 4B**), suggesting that the compound interacts with a cell surface molecule that is accessible independent of viral exposure.

**Fig 4:**
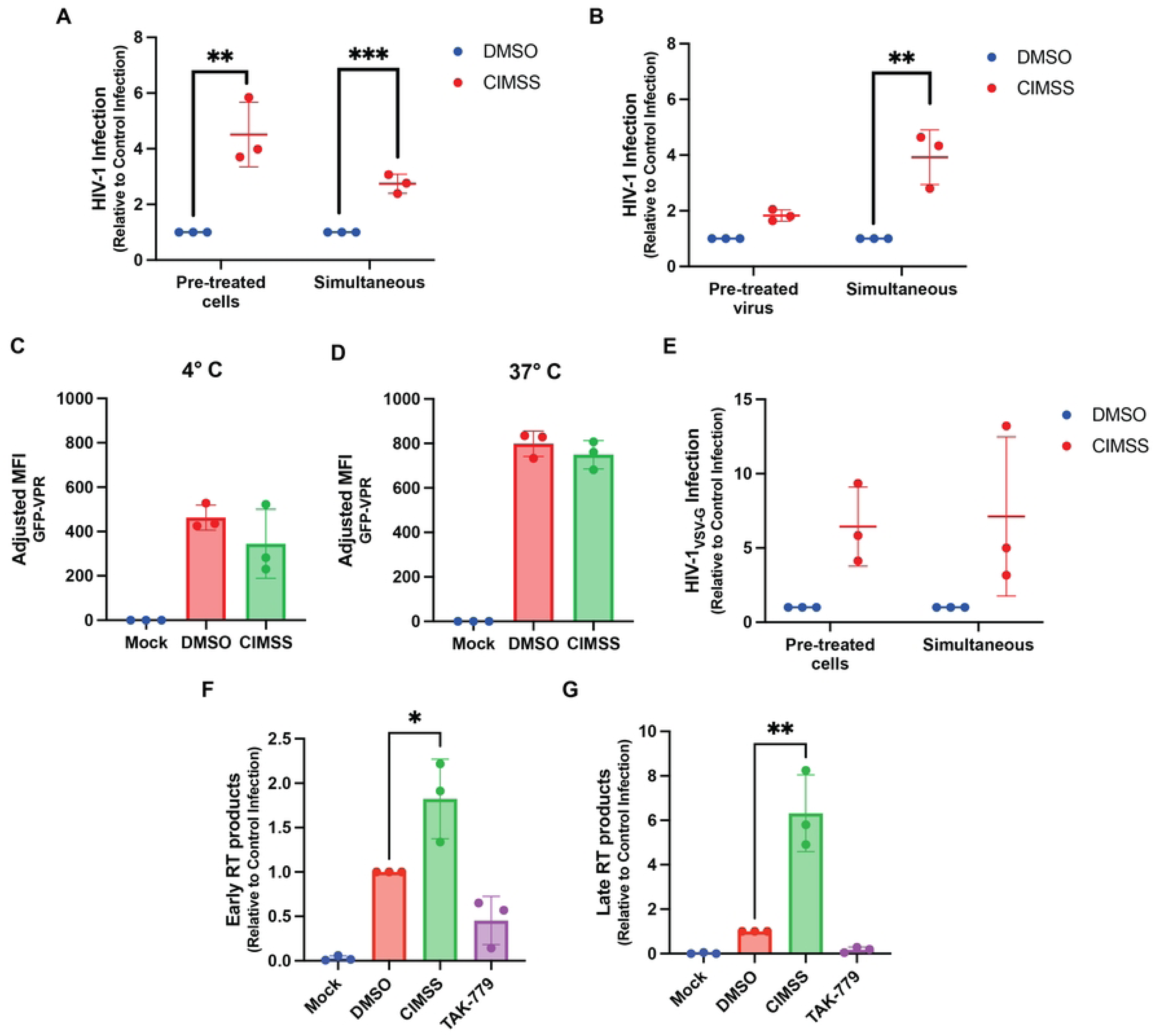
Alkyl-CIMSS acts on cells prior to viral exposure to promote the production of early and late gene transcripts. (A) TZM-bl cells were pre-treated with or DMSO vehicle control for 1 hour and then washed before infection with 1.5 ng p24 HIV-1_BaL_ or (B) concentrated HIV-1_BaL_ was treated with for 1 hour and then the virus-drug mixture diluted 40-fold (final alkyl-CIMSS concentration 1 μM) prior to infecting TZM-bl cells. In parallel, cells were exposed simultaneously to virus and compound. Cells were harvested 72 hpi and analyzed for luciferase activity. Results are presented as fold-change over infection in the presence of DMSO (n=3 independent experiments each conducted in triplicate). (C and D) Jurkat-CCR5 cells were treated with 50 μM alkyl-CIMSS or vehicle control and exposed to HIV-1 carrying a GFP-VPR fusion protein for 2 h either at 4° C or 37° C prior to being washed extensively. Cell associated virus was quantified by measuring GFP expression by flow cytometry and results are presented as the MFI GPF after subtracting background from mock infected cells (n=3 independent experiments each conducted in duplicate). (E) TZM-bl cells were pre-treated with 50 μM alkyl-CIMSS or vehicle control before or infected simultaneously with HIV-1_VSV-G_. Infection was monitored by assaying for luciferase activity 72 h (n=3 independent experiments each conducted in triplicate). (F and G) Jurkat-CCR5 cells were treated with 50 μM alkyl-CIMSS 10 μM TAK-779, or DMSO control before infection with 5 ng p24 HIV-1_BaL_. At 6 and 24 hpi, cells were harvested for DNA and quantification of early and late reverse transcription products, respectively (n=3 independent experiments each conducted in duplicate). In E-G, results are presented as mean ± SD fold-change over infection in the presence of DMSO and compared to the DMSO control by unpaired t-test; *P < 0.05, **P < 0.01, ***P <0.001.

To determine if alkyl-CIMSS promotes HIV-1 binding, entry, or acts post-entry, we took advantage of a virus carrying a Vpr-enhanced green fluorescence protein fusion protein (Vpr-EGFP), which is packaged into the virion during assembly, allowing for detection of bound and internalized virus. Jurkat-CCR5 cells were treated with 50 μM alkyl-CIMSS or control buffer for 4 hours and then exposed to the Vpr-EGFP viral construct for 2 hours at 4°C, a temperature only permissive for viral binding, washed, and bound virus detected by assaying for cell bound EGFP. Alternatively, after the 2 h viral binding period, the cells were shifted to 37°C for an additional 2 hours to allow viral entry. No differences in EGFP were detected in the absence or presence of alkyl-CIMSS (**Figs 4C and D**), indicating that viral attachment is independent of the CIMSS-dependent process. To further assess if alkyl-CIMSS enhanced HIV-1 infection post-entry, we tested the effects of the compound on a single-cycle VSV-G pseudotyped HIV-1 virus, which binds to cells primarily through interactions with the low density lipoprotein receptor (LDLR) and enters by endocytosis. There was a 6.4 ± 2.7-fold increase in luciferase production when TZM-bl cells were pretreated with alkyl-CIMSS or when the VSV-G pseudotyped virus and the compound were added simultaneously to cells (**Fig 4E**). Following entry, HIV-1 uses reverse transcriptase to produce early and late transcripts. To assess whether CIMSS promotes this step, Jurkat-CCR5 cells were again infected with HIV-1_BaL_ in the presence of alkyl-CIMSS or control DMSO buffer and DNA was isolated for quantification of early and late reverse transcription products. TAK-779 was included as an inhibitory control. Alkyl-CIMSS significantly increased the amount of early reverse transcription products 6 h post-infection and late reverse transcription products 24 h post-infection, whereas cells infected in the presence of TAK-779 had lower levels of both early and late transcription products, reflective of less virus entering the cell (**Figs 4F and 4G**). Taken together, these data suggest that alkyl-CIMSS interacts with a cell surface molecule accessible prior to viral exposure, which leads to an increased production of reverse transcriptase products independent of viral entry.

### HIV-1 enhancement is linked to upregulation of GPC1

To explore how alkyl-CIMSS affects the cells to promote HIV-1 infection, we performed bulk RNA-sequencing and whole cell proteomics on Jurkat-CCR5 cells treated with alkyl-CIMSS or DMSO control buffer with or without HIV-1 infection. The cells were lysed after 8 h, a timepoint selected based on the observation that there was an increase in early viral transcripts within this timeframe. Differentially expressed gene transcripts and proteins were defined by log2 fold-change (FC) > 0.5 and adjusted p-value < 0.05. Using these criteria, there were two candidates (one upregulated and one downregulated) that exhibited significant changes at both the RNA and protein level in alkyl-CIMSS-treated vs DMSO-treated HIV-1 uninfected cells (**Fig 5A and S2 Table**). GPC1 was identified as the transcript (FC=3.2, adj. p-value=1.0E-211) and protein (FC=2.8, adj. p-value=1.5E-5) most upregulated in response to alkyl-CIMSS in the uninfected and the second most upregulated protein in the HIV-1 infected cells (FC=2.1, adj. p-value=0.0015, **Fig 5B and S3 Table**). Notably, neither CD4 nor CCR5 expression were increased in response to alkyl-CIMSS (**S2 Table**).

**Fig 5:**
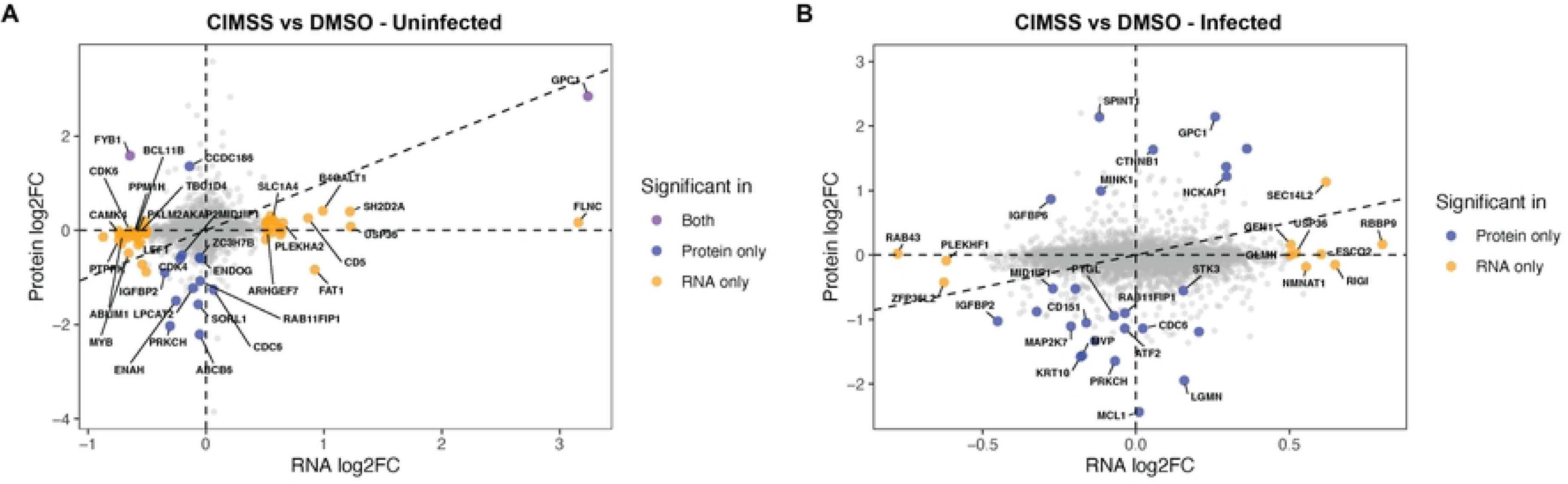
Proteomic and bulk RNA sequencing identifies changes in cells following exposure to alkyl-CIMSS. (A) Jurkat-CCR5 cells were treated with 50 μM alkyl-CIMSS or DMSO buffer or (B) subsequently infected with HIV. The cells were lysed after 8 hours and processed for proteomics and RNA-sequencing. The X-axis indicates the log2 fold chain (FC) in RNA and Y-axis indicates log2FC in protein expression. A transcript or protein was considered significant if the adjusted p-value was < 0.05 or log2FC > 0.5. Purple denotes significant changes at both RNA and protein level, blue denotes significance only at protein level, and yellow significance only at the RNA level.

We confirmed that alkyl-CIMSS increased GPC1 by directly quantifying gene and protein expression. Within 4 hours of exposure, alkyl-CIMSS treatment significantly increased *GPC1* RNA relative to control treated cells (7.2 ± 2.6-fold, p<0.05) (**Fig 6A**). Immunoblots revealed two GPC1 protein species, with a significant time-dependent increase in the lower (∼60kDa) band following alkyl-CIMSS treatment, which likely reflects newly synthesized protein lacking complete post-translational glycosylation (**Fig 6B**). There was also increase in GPC1 expression detected by flow cytometry on nonpermeabilized cells, consistent with GPC1 being a membrane protein (**Fig 6C**). In addition, increased GPC1 was detected in alkyl-CIMSS treated, HIV-infected PBMC (specifically in the CD3+/CD8^-^ subpopulation) but not in the control treated HIV-infected PBMCs (**Fig 6D**). Treatment of TMEM16F KO cells with alkyl-CIMSS also induced an increase in GPC1, which was associated with a significant increase in HIV-1 infection of the KO cells (p<0.05) (**Fig 6E and F).** However, infection remained less in the alkyl-CIMSS treated KO compared to control cells, indicating that the increase in GPC1 only partially overcame the absence of TMEM16F.

**Fig 6:**
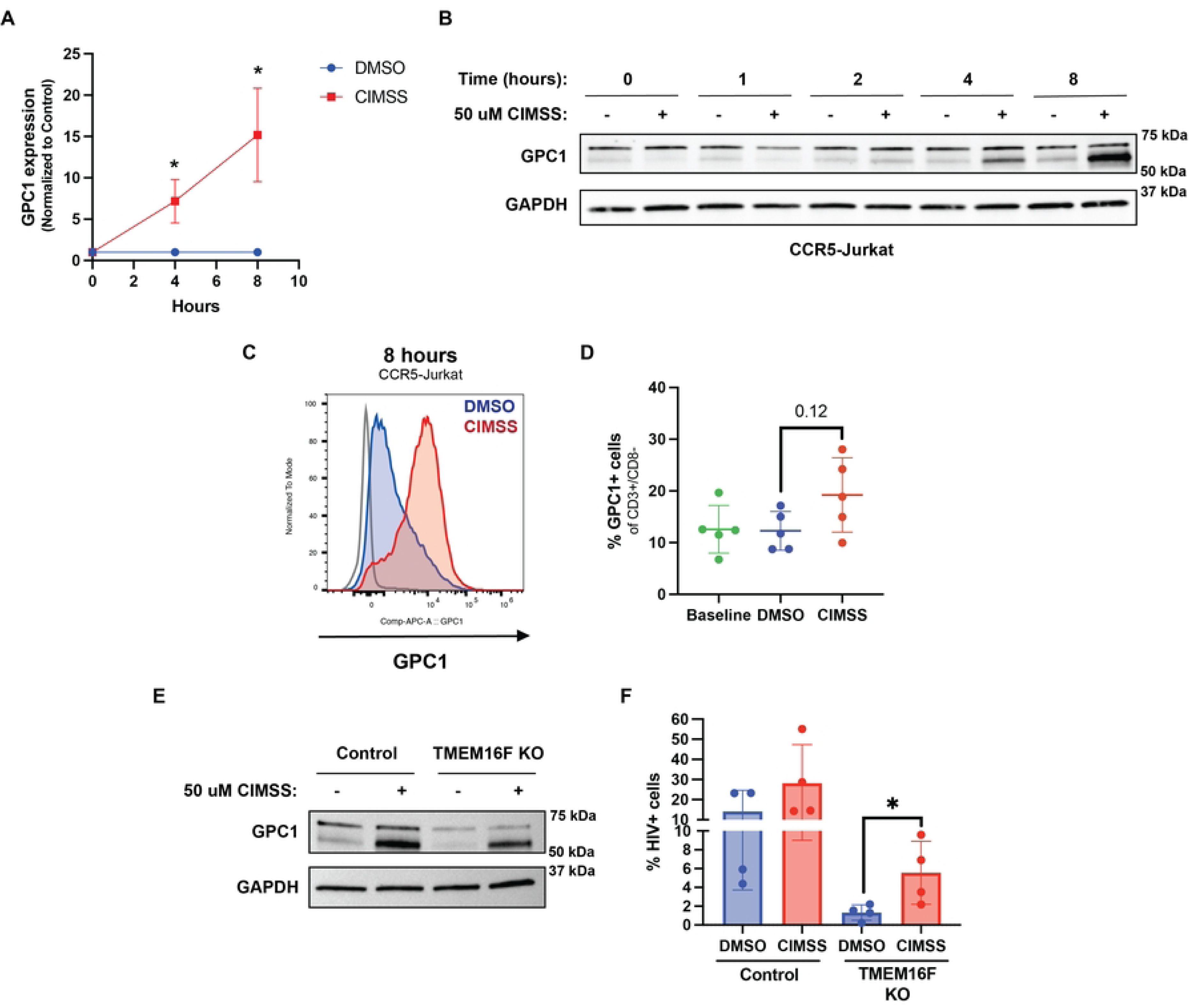
Kinetics of alkyl-CIMSS upregulation of GPC-1. (A) Jurkat-CCR5 cells or primary CD4+ T cells were treated with 50 μM alkyl-CIMSS or vehicle control for the indicated times and GPC-1 gene expression or (B) protein expression monitored by RT-qPCR (n=3) or (B) immunoblotting (n=2). (C) Jurkat-CCR5 (n=2 independent experiments) were treated as in A and GPC1 quantified at 8 hours by flow cytometry in nonpermeabilized cells. Histogram depicts the MFI of exofacial GPC1 on DMSO (blue) or alkyl-CIMSS (red) treated cells 8-hours post treatment. (D) PBMC cells (n=5) were treated with 50 uM alkyl-CIMSS or vehicle control and infected for 5 days. At 5 dpi, PBMCs were stained as in (C) for GPC1. PBMCs were also stained for CD3 and CD8. Results are presented as the percentage of GPC1+, CD3^+^CD8^-^ cells. (E) TMEM16F KO or CRISPR control cells were treated with alkyl-CIMSS or DMSO control for 8 hours. Cells were harvested and analyzed for GPC1 protein expression by western blotting or (F) infected with HIV-1_BaL_ and stained for intracellular p24 72 hpi (n = 4). Results are presented as mean ± SD and compared by unpaired (A, F) or paired (D) t-test; *P < 0.05.

### GPC1 enhances HIV-1 infection

To test whether the increase in GPC1 directly contributes to the enhanced HIV-1 infection observed in response to alkyl-CIMSS, Jurkat-CCR5 cells were transduced with a full-length GPC1 or control lentiviral construct expressing mCherry. The cells were then sorted and overexpression of GPC1 confirmed by flow cytometry for cell-surface GPC1 (**Fig 7A**). The transduced cells were infected with HIV-1 (5 ng p24), culture supernatants harvested at 72 hours and viral yields quantified by TZM-bl assay. There was a significant increase in HIV-1 production in GPC1-transduced cells compared to control cells (4.1 ± 0.3-fold, p<0.001) (**Fig 7B)** Conversely, the cells were transduced with an mCherry+ GPC1-silencing shRNA or control shRNA lentiviral construct. Cells were sorted and knockdown was assessed by flow cytometry, and the cells were infected with HIV-1. There was a modest but significant decrease in HIV-1 production compared to control cells (p<0.01) (**Fig 7C and D**). This was partially overcome when the cells were treated with alkyl-CIMSS, which also led to an increase in GPC-1 expression in the shRNA transduced cells (**Fig 7E and F**).

**Fig 7:**
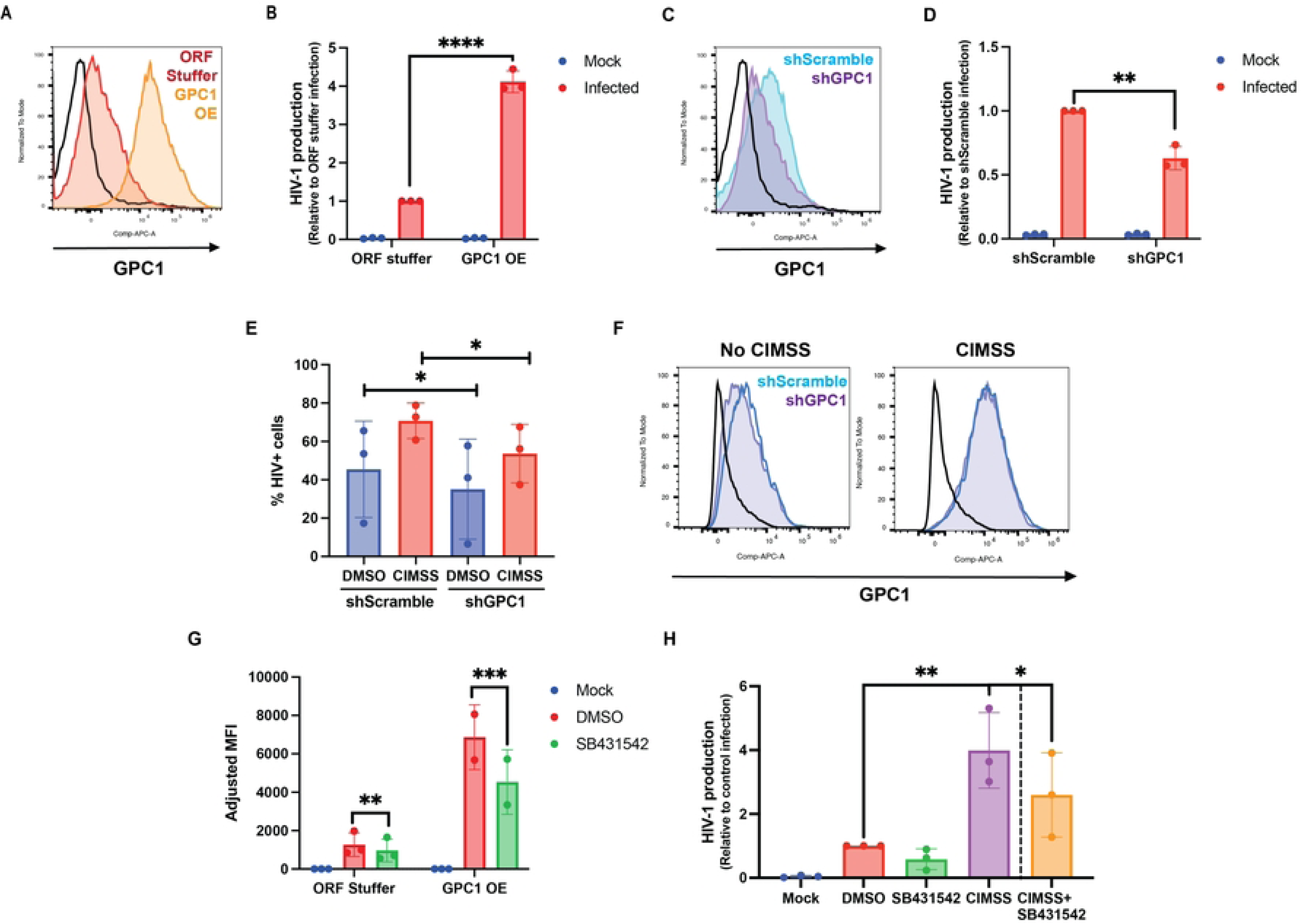
Glypican-1 enhances HIV infection and both alkyl-CIMSS and glypican-1 enhancement is reduced when cells are treated with an inhibitor of TGF-β. (A) Jurkat-CCR5 cells were transduced with a full-length glypican-1 overexpression (GPC1 OE) or control (ORF stuffer) construct. GPC1 overexpression was assessed by flow cytometry. (B) OE or control cells were infected with HIV-1_BaL_ and supernatant was collected 72 hpi for measurement of HIV-1 production by TZM-bl assay. Results are presented as fold-change over infection of control cells (n = 3). (C) Jurkat-CCR5 cells were transduced with a GPC1 shRNA (shGPC1) or scramble shRNA (shScramble) construct; silencing was assessed by flow cytometry. (D) shGPC1 or shScramble cells were infected with HIV-1_BaL_ and supernatant was collected 72 hpi for measurement of HIV-1 production by TZM-bl assay. Results are presented as fold-change over infection of control cells (n = 3). (E) shScramble or shGPC1 cells were treated with DMSO or alkyl-CIMSS and infected with HIV-1BaL. At 72 hpi, cells were stained for intracellular p24 or (F) exofacial GPC1 for measurement by flow cytometry. (G) GPC1 OE and control cells were infected with HIV-1_BaL_ in the presence of 1 μM SB431542 for 72 hours and then stained for intracellular p24. Adjusted MFI indicates p24 MFI of infected sample minus MFI of mock infected cells (n = 2). (H) Jurkat-CCR5 cells were treated with alkyl-CIMSS or DMSO control in the absence or presence of 1 μM SB431542 for 72 h. Infection was monitored by measuring HIV yields in culture supernatants in a TZM-bl assay. Results are presented mean ± SD fold change relative to infection in the controls (n=3). (B, D) are compared by unpaired and (E, G) paired t-test. In H, CIMSS and SB431542 are compared to DMSO by one-way ANOVA and alkyl-CIMSS+SB431542 is compared to alkyl-CIMSS by t-test (*p < 0.05, **p < 0.01, ***p <0.001, ****p<0.0001).

### Increased HIV-1 infection in response to alkyl-CIMSS is mediated in part by TGF-β

GPC1 regulates multiple cell signaling pathways including TGF-β, which has been previously shown to promote HIV-1 infection [13–15]. To explore whether TGF-β signaling contributed to the increase in HIV-1 infection associated with upregulation of GPC1, we treated GPC1 overexpressing cells or alkyl-CIMSS treated Jurkat-CCR5 cells with SB431542, a pharmacological inhibitor of TGF-β signaling [16]. SB431542 reduced

HIV-1 infection of both the GPC1 overexpressing (**Fig 7G**) and alkyl-CIMSS-treated Jurkat-CCR5 cells (**Fig 7H**). The drug also inhibited infection of the respective control cells, although infection of the controls was less than GPC1 overexpressing or alkyl-CIMSS-treated cells.

## Discussion

We took advantage of a cell-impermeable kinase inhibitor as a tool compound to test the hypothesis that HIV-1-mediated activation of TMEM16F induced exofacial movement of Akt as previously observed in response to HSV or SARS-CoV-2-mediated activation of PLSCR1. CIMSS (acyl and alkyl-CIMSS) blocked the phosphorylation of exofacial Akt and prevented HSV and SARS-CoV-2 entry. Surprisingly, we found that not only did alkyl-CIMSS fail to inhibit but rather enhanced HIV-1 infection. HIV-1, which was previously shown to activate a different phospholipid scramblase, TMEM16F, triggered the translocation of PS but not the exofacial movement of Akt. We confirmed a role for TMEM16F in HIV-1 but not HSV infection by generating TMEM16F KO Jurkat-CCR5 cells. The cells were significantly less susceptible to HIV-1 infection, but the KO had no discernible effect on susceptibility to HSV. These findings suggest that while both TMEM16F and PLSCR1 catalyze the translocation of membrane phospholipids, only PLSCR1 activation triggers the exofacial movement of Akt. Consistent with this notion, a recent study examined the TMEM16F-dependent remodeling of the plasma membrane comparing WT and TMEM16F KO Jurkat cells. Activation of TMEM16F by ionomycin resulted in changes in the extracellularly exposed proteome primarily associated with antigen presentation and T cell receptor signaling but no increase in Akt was detected [17].

In addition to the finding that Akt was only detected by flow cytometry in nonpermeabilized cells following HSV but not HIV-1 exposure, the observation that alkyl-CIMSS enhanced HIV-1 infection when the cells were pretreated with the tool compound then washed suggests that a cell surface protein other than Akt, possibly a different kinase, is the direct target of alkyl-CIMSS. Since alkyl-CIMSS also enhanced HIV-1 infection of TMEM16F KO cells, this target molecule must be present on the outer leaflet of the plasma membrane independent of scramblase activation.

Whole cell proteomic and transcriptomic studies of alkyl-CIMSS treated cells identified increased expression of GPC1 as a candidate for mediating the enhancement of HIV-1 infection. This hypothesis was confirmed by showing that overexpression of GPC1 also increased HIV-1 infection. The enhancing effects of GPC1 were independent of TMEM16F activation as alkyl-CIMSS upregulated GPC1 and partially restored the susceptibility of TMEM16F KO cells to HIV-1 infection. Prior studies have shown that TMEM16F primarily acts by promoting HIV-1 entry [8]. In contrast, alkyl-CIMSS promoted HIV-1 infection post-entry processes as evidenced by our observations that there was no increase in bound or internalized viral particles when CIMSS-treated cells were exposed to a Vpr-EGFP reporter virus and enhancement was preserved when cells were infected with an HIV-1 virus pseudotyped with VSV-G, which binds and enters by a completely different pathway.

We speculate that alkyl-CIMSS and the associated upregulation of GPC1 might promote the generation of reverse transcriptase products through TGF-β signaling (**Fig 8**). This notion was suggested by prior studies showing that GPC1 can signal through TGF-β [18–20], and TGF-β has been shown to promote HIV-1 infection by complex mechanisms, including upregulation of CCR5 and enhanced HIV-1 transcription [14, 15]. Addition of a pharmacologic inhibitor of TGF-β signaling reduced, but did not abolish, the enhancing effects of alkyl-CIMSS and GPC1 overexpression. We speculate that GPC1, which also functions as a co-receptor for a variety of signaling pathways in addition to TGF-β, including VEGF, FGF, and Wnt [21, 22], may act through multiple pathways to promote the generation of HIV-1 reverse transcriptase products.

**Fig 8:**
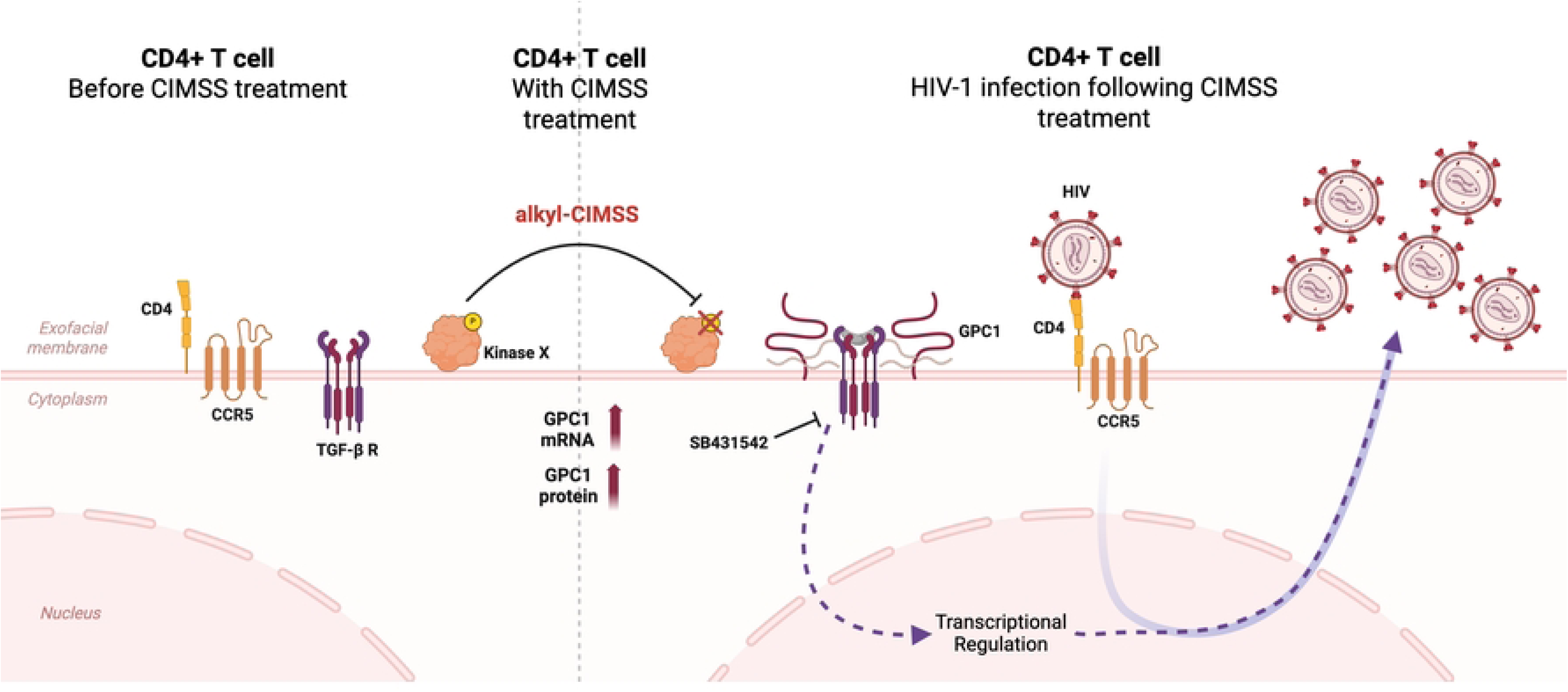
Model of Alkyl-CIMSS-triggered glypican-1 upregulation. CD4+ T cells have a basal expression of CD4, CCR5 and TGF-β receptors at the plasma membrane, rendering the cells susceptible to HIV. Akyl-CIMSS blocks the phosphorylation of exofacial kinase(s), which promotes the upregulation of glycican-1. Glypican-1, in turn, activates TGF-β and other signaling pathways to promote the generation of early and late HIV reverse transcriptase products. The TGF-β inhibitor, SB431542, partially abrogates the enhancing effects of glypican upregulation.

A limitation of these studies is that we have not yet identified the direct target of alkyl-CIMSS on T cells and how this then leads to upregulation of glypican-1. The target must be present in TZM-bl, Jurkat-CCR5, primary CD4+ T cells as well as TMEM16F knockout Jurkat-CCR5 cells since alkyl-CIMSS upregulated glypican in each of these cell types. Over 4000 extracellularly exposed proteins were detected in resting Jurkat cells and all but 465 were also detected in TMEM16F KO cells. These included many canonically cytosolic kinases such as PKCα, CDK1, CDK2 that may be linked to the upregulation of GPC1 and are subjects for future studies [17].

Despite this limitation, results herein demonstrate that cell-impermeable kinase inhibitors represent critical tools to study previously underappreciated “outside-in” signaling pathways. These tool compounds enabled our earlier studies, and confirmed here, with TZM-bl and Jurkat T cells, that HSV triggers the exofacial movement and subsequent phosphorylation of Akt to facilitate viral entry. In contrast, the same cell-impermeable pan-kinase inhibitor targets a different and as of yet unidentified molecule or molecules present on the cell surface, independent of viral exposure, which upregulate GPC1 to enhance HIV-1 infection. Defining how the exofacial proteome drives outside-in signaling to promote HIV-1 infection may enable the development of novel antiviral drugs that target these pathways. The tool compounds also provide the opportunity to explore not only how viruses but also normal physiological processes signal through the exofacial proteome.

## Materials and methods

### Cells

TZM-bl cells were obtained through the NIH HIV Reagent Program, Division of AIDS, NIAID, NIH. Jurkat-Tat-CCR5 (Jurkat-CCR5) cells were provided by Quentin Sattentau (Sir William Dunn School of Pathology, University of Oxford, Oxford, UK). PMBC were obtained from HIV-uninfected donors (New York Blood Center) and CD4+ T cells were isolated using negative selection (EasySep Human CD4+ T cell Isolation Kit, 19052, STEMCELL Technologies) and cultured in RPMI medium with 50 U/mL recombinant IL-2 (#202-IL, R&D Systems). Primary CD4+ T cells were activated for 72 hours with 25 uL/mL ImmunoCult Human CD3/CD28 T cell activator (10971, STEMCELL Technologies). Whole PBMCs were cultured in complete RPMI medium.

TMEM16F knockout Jurkat CCR5 cells were produced using Dharmacon’s Edit-R CRISPR-Cas9 All-in-One lentiviral technology. Briefly, Jurkat-Tat-CCR5 cells were transduced with lentivirus containing Cas9, sgRNA targeting TMEM16F, and an EGFP selection marker at a multiplicity of infection (MOI) of 0.3 transducing units (TU) per cell. At 72 hours post transduction, single clones were isolated for outgrowth using the BD FACSARIA. Gene expression and knockdown were validated by RT-qPCR and flow cytometry. Controls were generated by transducing cells with a non-targeting control CRISPR/Cas9 construct and following the same cloning strategy.

Jurkat-CCR5 cells overexpressing GPC1 were generated by lentiviral transduction using lentiviral vectors constructed by VectorBuilder. These pseudoviruses were produced by co-transfecting HEK293T cells in 10 cm^2^ dishes with 3 ug of plasmid DNA, 0.3 ug pMD2.G vector expressing VSV-G, and 2.5 ug pCMVΔ8.2. FuGENE HD Transfection Reagent (Promega, E2311) was used at a 3:1 ratio with plasmid DNA. pCMVR8.74 (Addgene plasmid #22036) and pMD2.G (Addgene plasmid # 12259) were gifts from Didier Trono. Pseudoviruses were added to Jurkat-CCR5 cells and incubated at 37°C for 3 days. After 3 days, mCherry bright live cells were sorted using BD FACSARIA for outgrowth. Gene expression and knockdown were validated by RT-qPCR and immunoblots. Control cells were stably transfected with an empty open reading frame and mCherry. A similar approach was used to generate Jurkat-CCR5 GPC1 knockdown cells by transducing the cells with lentiviruses expressing shRNA specific to GPC1 and mCherry or a shSramble shRNA and mCherry.

### Viruses and Pseudoviruses

HIV-1_BaL_ (R5-utilizing strain) and HIV-1_iiiB_ (X4-utilizing strain) were acquired through the NIH AIDS Reagent Program, Division of AIDS, National Institute of Allergy and Infectious Diseases (NIAID), NIH, and grown on Jurkat (X4) or Jurkat-CCR5 (R5) cells for 12 days and stored at -80°C after filtration through a 0.2μM filter. Viral stocks were quantified by p24 ELISA (R&D Systems, DY7360). Fluorescently labeled HIV-1 and pseudoviruses were produced by transfecting HEK293T in T150s using the ViaFect Transfection Reagent (Promega, E4981). EGFP labeled HIV-1 was produced by co-transfecting 60 ug pNL4-3_BaL_ p21-85 (NIH HIV Reagent Program, ARP-11441, contributed by Dr. Bruce Chesebro), 20 ug pEGFP-Vpr (NIH HIV Reagent Program, ARP-11386, contributed by Dr. Warner C. Greene), and 10 ug of pAdvantage (Promega, E1711). VSV-G pseudotyped HIV-1 was produced by co-transfecting 60 ug of pNL4-3Δenv (BEI Resources, NIAID, NIH, HRP-20281), 20 ug VSV-G expression vector (NIH HIV Reagent Program, ARP-4693, contributed by Dr. Lung-Ji Chang), and 10 ug pAdvantage. All HIV-based viruses were quantified by p24 ELISA (R&D Systems, DY7360). HSV-2 (G) and HSV-1(F-GS2822), which encodes red fluorescent protein fused to the N terminus of VP26 [23], were grown and titered on Vero cells as previously described [24, 25].

### Synthesis of alkyl-CIMSS

Alkyl-CIMSS was synthesized in a similar manner to that used to synthesize CIMSS previously in our group [6, 26]. Briefly, staurosporine was alkylated with methyl bromoacetate and saponified to afford known acid **2** [27–30]. This acid was engaged in a HATU-mediated amide coupling with known sulfonate-containing amine **S3** [6, 26] which, after neutralization, afforded alkyl-CIMSS **S5** in its zwitterionic form. Full details of the synthesis of alkyl-CIMSS is found in S1 Appendix.

### Flow cytometry to assess for phosphatidylserine and protein expression

Jurkat-CCR5 or TZM-bl cells were treated with either HIV-1_BaL_ (∼10 ng p24/5x10^5^ cells) or HSV-2(G) (MOI 10) for 30 minutes or control buffer. The cells were then washed and stained with zombie NIR fixable viability dye (BioLegend, #423105). PS was detected using mouse anti-PS IgG, clone 1H6 (0.2 ug/mL, MilliporeSigma, #05-719) followed by goat-anti-mouse PE (1:200; Invitrogen, #31861). Akt was detected using anti-Akt (pan) from Cell Signaling (1:50, #5186). GPC1 was detected using anti-GPC1 (1:100, Invitrogen, PA5-28055) followed by goat-anti-rabbit APC (1:500, Invitrogen, A10931). HIV p24 was detected in permeabilized cells with anti-HIV1 p24 (NIH HIV Reagent Program, ARP-13449, contributed by DAIDS/NIAID). Cells were fixed in 2% paraformaldehyde. Permeabilized cells generated using Cytofix/Cytoperm (BD, #554714) were used as staining controls. Samples were acquired on a Cytek Aurora flow cytometer (Cytek Biosciences) and analyzed with FlowJo (FlowJo LLC).

### Viral infection assays

Cells were infected with HSV-2(G) or HIV-1 (5 ng p24 unless otherwise indicated for 2 hours at 37°C. For nonadherent cells (Jurkat-CCR5 and PBMC), the infected cells were washed and transferred to 24-well plates with 1 mL of complete media for 72 hours. At 72 hours, supernatants were harvested and stored at -80°C for TZM-bl assays or the infected cells were permeabilized and assayed for p24 expression by flow cytometry. Alternatively, cells were lysed for RNA extraction and *ltr* expression. TZM-bl assays were conducted as previously described [31, 32]. After 48 h incubation, the cells were lysed in passive lysis buffer (Promega, #E1941) and luciferase activity measured using the Promega assay system (Promega, #E1501).

PBMCs (2x10^6^ cells) were inoculated with 50 ng p24 in the presence of alkyl-CIMSS or DMSO. Every two days, half of the media was exchanged to resupplement vehicle or drug. At 5 days post infection, media was collected and analyzed for p24 content by ELISA (R&D Systems, DY7360) or viral yield by TZM-bl assay. Values below the limit of detection were adjusted to 0.

Cells were infected with HSV-1(G) at MOI of 10 pfu/cell and infection monitored by flow cytometry for immediate early gene ICP0 or RT-qPCR as described below.

### Pharmacologic inhibitors

TAK-779 (MedChemExpress, #HY-13406), AMD3100 (MilliporeSigma, 239825), CaCCinh-A01 (MedChemExpress, #HY-100611), SB431542 (StemCell, #72232), and DMSO (Cell Signaling, #12611) as a carrier control were added 5 min before application of virus. Concentrations used were determined non-toxic before use via MTS assay (Promega, G3582).

### RNA extraction and real-time quantitative reverse transcription polymerase chain reaction (RT-qPCR)

Total RNA was extracted, cDNA was synthesized, and reverse-transcription polymerase chain reaction amplification was performed as described elsewhere [17]. Primers/probes used were as follows: HSV *ICP0* (forward: 5′-GGTCACGCCCACTATCAGGTA-3′; reverse: 5′-CCTGCACCCCTTCTGCAT-3′; probe: 5′-FAM-CAACGGAATCCAGGTCTTCATGCACG-TAMRA-3′); HIV *LTR* (forward: 5′-CACACAAGGCTACTTCCCTGA-3′; reverse: 5′-TCTCTGGCTCAACTGGTACTAGCTT-3′; probe: 5′-FAM-AGAACTACACACCAGGGCCAGGGATCAG-TAMRA-3′); TMEM16F (*ANO6*) (ThermoFisher Scientific, Hs03805835_m1); *GPC1 (*ThermoFisher Scientific, Hs00892476_m1); *RPLP0* (ThermoFisher Scientific, 4326314E), and *GAPDH* (ThermoFisher Scientific, Hs02786624_g1). Targets were amplified in 10 μL reactions using TaqMan Gene Expression Master Mix (Applied Biosciences, #4369016) in a QuantStudio 7 Flex Real-Time PCR System (Thermo Fisher Scientific). Data were analyzed using QuantStudio software. Quantification was normalized against the housekeeping gene *RPLP0* (T cell lines) or *GAPDH* (TZM-bl cells) in the same RNA extracts; relative gene expression was calculated using the 2^−ΔΔCt^ method.

### HIV-1 binding and entry assays

Cells were pre-treated with 50 μM CIMSS or DMSO for 4 hours at 37°C. Following this incubation, HIV_NL4.3BaL/VPR-EGFP_ (50 ng p24) was applied to the cells and allowed to incubate for 2 hours at either 4°C (binding) or 37°C (entry). The cells were washed to remove any unbound virus and fixed for analysis by flow cytometry.

### Quantification of early and late HIV-1 reverse transcripts

Jurkat-CCR5 cells (10^5^) were pre-treated with compounds for 15 minutes at 37°C before infection with 5 ng p24 HIV-1BaL for 2 hours. The cells were then washed thrice with media to remove any unbound virus and resuspended in complete media. At the indicated times, cells were washed with PBS and harvested for DNA extraction using the DNeasy Blood & Tissue Kit (Qiagen, 69506). 20 ng genomic DNA was used per 10 μL real-time PCR reaction containing 2.5 nanomoles of probe and 5 nanomoles of each primer with TaqMan Gene Expression Master Mix (AppliedBiosciences, #4369016). Primer/probes were: Early Reverse Transcription product (ERT; forward: 5′-GTGCCCGTCTGTTGTGTGAC-3′; reverse: 5′-GGCGCCACTGCTAGAGATTT-3′; probe: 5′-FAM-CTAGAGATCCCTCAGACCCTTTTAGTCAGTGTGG-TAMRA-3′) and Late Reverse Transcription product (LRT; forward: 5′-TGTGTGCCCGTCTGTTGTGT-3′; reverse: 5′-GAGTCCTGCGTCGAGAGAGC-3′; probe: 5′-FAM-CAGTGGCGCCCGAACAGGGA-TAMRA-3′). Standard curves were generated by plotting Ct values against dilutions of pNL4-3 (NIH HIV Reagent Program, ARP-114, contributed by Dr. M. Martin) at known concentrations.

### Whole cell proteomics

Jurkat-CCR5 cells were treated with 50 μM alkyl-CIMSS or 0.5% DMSO as a vehicle control before being either mock-infected or infected with HIV-1_BaL_ for 8 hours. At 8 hours, cells suspensions were transferred to conical tubes, pelleted and resuspended in urea lysis buffer (8M urea, 50 mM NH4HCO3, 150 mM NaCl. For sample processing, Tris-(2-carboxyethyl) phosphine (TCEP) was added to a final concentration of 4 mM. The DNA was sheared via probe sonication, on ice, at 20% amplitude for 20 sec, followed by 10 sec of rest for a total of three times. After sonication, protein concentration was determined using BCA assay (Thermo, 23225). Then, Iodoacetamide (IAA) was added to each sample to a final concentration of 10 mM, and samples were incubated in the dark at room temperature (RT) for 30 minutes. Excess IAA was quenched by the addition of dithiothreitol (DTT) to 10 mM, followed by incubation in the dark at RT for 30 minutes. Samples were then diluted with 0.1 M NH4HCO3 (pH = 8.0) to a final urea concentration of 2 M. Trypsin (Gold mass spectrometry grade, Promega) was added at a 1:100 (enzyme:protein w:w) ratio and digested overnight at 37 °C with rotation. Following digestion, 10% trifluoroacetic acid (TFA) was added to each sample to a final pH ∼2. Samples were desalted under vacuum using Sep Pak tC18 cartridges (Waters). Each cartridge was activated with 1 mL 80% acetonitrile (ACN)/0.1% TFA, and equilibrated with 3 x 1 mL of 0.1% TFA. Following sample loading, cartridges were washed with 4 x 1 mL of 0.1% TFA, and samples were eluted with 4 x 0.5 mL 50% ACN/0.25% formic acid (FA). All samples were analyzed on an Orbitrap Eclipse mass spectrometry system equipped with an Easy nLC 1200 ultra-high pressure liquid chromatography system interfaced via a Nanospray Flex nanoelectrospray source (Thermo Fisher Scientific). Samples were injected onto a fritted fused silica capillary (30 cm × 75 μm inner diameter with a 15 μm tip, CoAnn Technologies) packed with ReprosilPur C18-AQ 1.9 μm particles (Dr. Maisch GmbH). Buffer A consisted of 0.1% formic acid in water, and buffer B consisted of 0.1% formic acid in 80% acetonitrile. Peptides were separated by an organic gradient from 5% to 35% mobile buffer B over 120 min, followed by an increase to 100% B over 10 min at a flow rate of 300 nL/min. Analytical columns were equilibrated with 3 μL of buffer A.

Data were acquired in a data-independent analysis (DIA) manner. A full scan was collected at 60,000 resolving power over a scan range of 390-1010 m/z, an instrument controlled AGC target, an RF lens setting of 30%, and an instrument controlled maximum injection time, followed by DIA scans using 8 m/z isolation windows over 400-1000 m/z at a normalized HCD collision energy of 28%.

The DIA-NN algorithm was used to identify peptides/proteins and extract intensity information from DIA data [33]. A library-free FASTA digest was used for library generation using the Homo sapiens UniProt reference proteome (downloaded on August 23, 2023) and HIV-1 (strain NL4-3) protein sequences. Search setting were the algorithm defaults: the protease setting was Trypsin/P, maximum missed cleavages was 1, fixed modifications for N-terminal methionine excision and cysteine carbamidomethylation were included, the peptide length 7-30 amino acids, the precursor charge range was 1-4, the precursor *m/z* range was 300-1800, the fragment ion *m/z* range was 200-1800, and the precursor false discovery rates was set to 1%.

Statistical analysis of proteomics data was conducted utilizing the MSstats package in R [34]. All data were normalized by equalizing median intensities, the summary method was Tukey’s median polish, the maximum quantile for deciding censored missing values was 0.999, and only the top 25 most abundant features per protein were included in modeling.

### Bulk RNA-Sequencing

Jurkat-CCR5 cells were treated with 50 μM alkyl-CIMSS or 0.5% DMSO as a vehicle control and then infected with HIV-1_BaL_ or control media for 8 hours. The cells were then lysed and RNA extracted using the RNeasy Plus Mini kit (Qiagen, #74134). Isolated RNA was sent to Azenta Life Sciences (New Jersey, USA) for library preparation and sequencing on the Illumina NovaSeq platform using 150 bp paired-end reads to generate 30 million read pairs per sample. Raw sequence data were processed using the nf-core RNA-Seq pipeline (version 3.18.0) [35, 36]. Reads were aligned to the GRCh38 human reference genome using the STAR aligner and gene quantification was performed against the GENCODE v42 primary assembly annotation with RSEM to obtain gene expression counts [37, 38]. Expressed genes were determined using zFPKM [39]. Normalization and differential expression analysis were performed using DESeq2 [40]. Gene set enrichment analysis was performed using clusterProfiler and GSVA [41, 42].

### Western blots

Total cell lysates were prepared from equal cell numbers using the Radio-immunoprecipitation Assay (RIPA) buffer system (SantaCruz, sc-24948) containing 1X phosphatase and protease inhibitor cocktail (ThermoFisher, 78440). SDS-PAGE gel electrophoresis was performed on a gradient polyacrylamide gel (Biorad, 5671083) for 80 minutes at 120 V and transferred on PVDF (Biorad, 1704157) using the Turboblot (Biorad, 1704150). Western blots were visualized and scanned using the ChemiDoc imaging system (Biorad). Protein loading was compared by staining with an antibody for GAPDH (1:1000, CellSignaling, 2118).

### Statistical analyses and data sharing

Statistical analyses of RT-qPCR data were performed using log10-transformed values including where data are presented as non-transformed values. Data were tested for normality and analyzed using parametric (normally distributed) or non-parametric (non-normally distributed) tests. P-value < 0.05 was considered significant. Analyses were performed using GraphPad Prism version 10.5.0 software (GraphPad Software Inc. San Diego, CA).

The input data and scripts used to generate the figures and numbers reported in this manuscript were deposited to the github repository: https://github.com/cutleraging/2025-Kelsey-Vinzant

## Acknowledgements

Flow cytometry studies were carried out using resources of FACS Core Facility of the Einstein Cancer Center, which is supported by NIH/NCI Cancer Center Service Grant P30 CA13330 and by Shared Instrumentation grants S10OD026833 and S10OD032169.

## Supporting information

**S1 Fig. Alkyl-CIMSS is non-toxic across cell types and does not trigger caspase activation.**

(A) Model of alkyl-CIMSS. (B) CCR5-Jurkat cells (∼50% confluence) were cultured in media containing increasing concentrations of Alkyl-CIMSS (12.5–50 μM) or the equivalent highest concentration of DMSO (0.5%) and cell proliferation and viability quantified after 24 h (n=3 independent experiments conducted in triplicate). (C) CCR5-Jurkat cells treated with either DMSO (blue), 50 uM Alkyl-CIMSS (orange), or 10 uM staurosporine (red) for 6 hours. Representative histogram (N=2) depicts poly-caspase activation as measured by flow cytometry.

**S2 Fig. Synthesis of Alkyl-CIMSS.**

Staurosporine was alkylated with methyl bromoacetate and saponified to afford known acid **2**. This acid was engaged in a HATU-mediated amide coupling with known sulfonate-containing amine **S3** which, after neutralization, afforded alkyl-CIMSS **S5** in its zwitterionic form.

**S3 Fig. Gating strategy used for the flow cytometry analysis of HIV-1BaL infection of CCR5-Jurkat cells.**

(A) Single cells were selected by gating SSC-A vs SSC-H (B). (C) Live vs dead cells were determined with the Zombie NIR fixable viability kit. (D) Cells were further analyzed for intracellular p24 expression.

**S4 Fig. Transcriptomics quality control.**

(A) Density plots Log_2_(FPKM) of each sample showing unfiltered distribution (blue) and the Gaussian fit to the Log_2_(FPKM) data, which represents the filtering threshold used to identify expressed genes used in downstream analyses. (B) Number of genes detected at TPM > 1 in each sample. (C) Distribution of DESeq2 normalized Log2(counts + 1).

(D) Principle component analysis (PCA) of all samples showing first 2 PCs.

**S1 Table. Biochemical characterization of Alkyl-CIMSS.**

(A) Permeability of Alkyl-CIMSS as determined by MDCK-MDR1 assay. (B) Enzyme activity in the presence of Alkyl-CIMSS.

**S2 Table. Transcriptomic and proteomic datasets from cells treated with Alkyl-CIMSS in the absence of HIV-1.**

**S3 Table. Transcriptomic and proteomic datasets from cells treated with Alkyl-CIMSS in the presence of HIV-1.**

